# Depletion of LONP2 unmasks differential requirements for peroxisomal function between cell types and in cholesterol metabolism

**DOI:** 10.1101/2023.03.15.532715

**Authors:** Akihiro Yamashita, Olesia Ignatenko, Mai Nguyen, Raphaëlle Lambert, Kathleen Watt, Caroline Daneault, Isabelle Robillard-Frayne, Ivan Topisirovic, Christine Des Rosiers, Heidi M. McBride

## Abstract

Peroxisomes play a central role in tuning metabolic and signaling programs in a tissue- and cell type-specific manner. However, the mechanisms by which the status of peroxisomes is communicated and integrated into cellular signaling pathways is not yet understood. Herein, we report the cellular responses to acute peroxisomal proteotoxic stress upon silencing the peroxisomal protease/chaperone LONP2. Depletion of LONP2 triggered accumulation of its substrates, alterations in peroxisome size and numbers, and luminal protein import failure. Gene expression changes and lipidomic analysis revealed striking cell specific differences in the response to siLONP2. Specific to COS-7 cells was a strong activation of the integrated stress response (ISR) and upregulation of ribosomal biogenesis gene expression levels. Common changes between COS-7 and U2OS cell lines included repression of the retinoic acid signaling pathway, and upregulation of sphingolipids. Cholesterol accumulated in the endomembrane compartments in both cell lines, consistent with evidence that peroxisomes are required for cholesterol flux out of late endosomes. These unexpected consequences of peroxisomal stress provide an important insight for our understanding of the tissue-specific responses seen in peroxisomal disorders.

## Introduction

Peroxisomes are ubiquitous eukaryotic organelles that are central to regulation of lipid metabolism, redox regulation and signaling (Wanders et al., 2023). The precise function of peroxisomes is coupled to cellular demand and the metabolic wiring within a given tissue (He et al., 2021). Peroxisomal metabolic pathways include catalase-mediated reduction of peroxide to water, and beta-oxidation of very long chain or branched chain fatty acids. Once fatty acid chains are reduced to shorter lengths, they are transferred to mitochondria for completion of beta-oxidation (Wanders et al., 2023). In liver, peroxisomes function with mitochondria and the endoplasmic reticulum (ER) to generate bile acids, a process that can utilize up to 90% of newly synthesized cholesterol in rodents (Vaz and Ferdinandusse, 2017). In the brain, peroxisomal factors are enriched in astrocytes compared to neurons, and cultured astrocytes but not neurons demonstrate beta-oxidation activity (Eraso-Pichot et al., 2018; Ioannou et al., 2019; McGee et al., 1992; Russo et al., 2021). In oligodendrocytes, a primary function of peroxisomes is the generation of plasmalogen ether lipids, as these lipids synthesized in peroxisomes and ER are essential for myelin formation by these cells (Kassmann et al., 2007). These examples highlight the fact that the mass metabolic flux through peroxisomes varies greatly depending on the tissue and need. Pathologies arise when peroxisomal function is altered by pathogenic variants. Peroxisomal biogenesis diseases are the most severe, manifesting with multisystemic developmental pathologies, growth failure, and early lethality (Wanders et al., 2023). Peroxisomal dysfunction is also associated with ageing, cancer, and neurodegenerative diseases, likely contributing to the pathogenesis of common diseases (Walker et al., 2018; Zalckvar and Schuldiner, 2022).

A key open question is how peroxisomes may communicate their status through stress responses and retrograde signaling pathways. New approaches to monitor transcriptional, lipidomic and proteomic changes in conditions of induced peroxisomal stress, primarily through the targeted loss of proteins essential for peroxisomal import, shed light on generalised organellar dysfunction. While global cellular stress responses are elicited in these conditions, there is a surprising variability in the details of these responses depending on the specific import component under investigation, the cell or tissue type and the model organism used (Mast et al., 2011; Rackles et al., 2021; Wangler et al., 2017). This hints that peroxisomal stress may not always lead to comparable stress responses, likely reflecting distinct functions of peroxisomal import components and their clients.

The loss of peroxisome import provides an excellent experimental approach to reduce or block peroxisomal function. In this study, we took a different approach to specifically monitor the stress response driven by an accumulation of unfolded proteins within the peroxisomal lumen. Currently, lon peptidase 2 (LONP2) is the only known peroxisomal protease that is conserved in plants, fungi and mammals (Goto-Yamada et al., 2014). LONP2 is the product of an early gene duplication of mitochondrial LONP1, which was retained from its bacterial origin (Gibellini et al., 2020). Lon peptidases act as both a chaperone and an ATP dependent protease responsible for the degradation and turnover of oxidized proteins in bacteria, mitochondria, peroxisomes and chloroplasts (Tsitsekian et al., 2019; Wlodawer et al., 2022). Studies have directly investigated the biochemistry and substrates of peroxisomal LONP2, where the protease TYSND1 is one substrate identified in mammalian cells (Okumoto et al., 2011), and other clients including catalase have been identified in the fungi *P. chrysogenum* and *H. polymorpha* (Aksam et al., 2007; Bartoszewska et al., 2012). In addition to LONP2 protease activity, its chaperone function was illustrated by the development of significant peroxisomal protein aggregates upon loss of LONP2 in fungi, thereby demonstrating that loss of LONP2 induced proteotoxic stress (Aksam et al., 2007; Bartoszewska et al., 2012). While the function of LONP2 as a key regulator of peroxisomal protein turnover and homeostasis is widely accepted, the functional consequences of these processes have not been explored in mammalian cells (Pomatto et al., 2017). To map the peroxisomal proteotoxic stress response that could be independent from a total loss of protein import, we took an siRNA approach to acutely silence expression of LONP2 in two distinct cell lines, COS-7 (African green monkey kidney cell line transformed with SV40) and the human osteosarcoma cell line U2OS. Imaging, RNAseq and lipidomic data reveal new findings that couple peroxisomal homeostasis to cholesterol trafficking, retinoic acid signaling, ribosome biogenesis and the integrated stress response.

## Results

### Depletion of peroxisomal LONP2 caused peroxisomal dysfunction in COS-7 cells and U2OS cells

To induce acute peroxisome-specific damage, we silenced LONP2 over 6 days in COS-7 and U2OS cells. Immunoblotting of *LONP2*-silenced cells after 6 days revealed an accumulation of the autocleaved products of the protease TYSND1, products established as LONP2 substrates (Okumoto et al., 2011), indicating that the efficiency of proteolytic processing inside peroxisomes was impaired (**Fig. 1a and c**). Analysis of peroxisomal membrane protein PMP70 or luminal protein thiolase (ACAA1) did not reveal a reduction in either of the protein levels, indicating that the peroxisomal mass was generally unchanged (**Fig. 1a and c**). Confocal microscopy demonstrated that peroxisomes were less abundant, but individual peroxisomes were elongated and enlarged in both cell lines (**Fig. 1b and d**).

**Figure 1.**
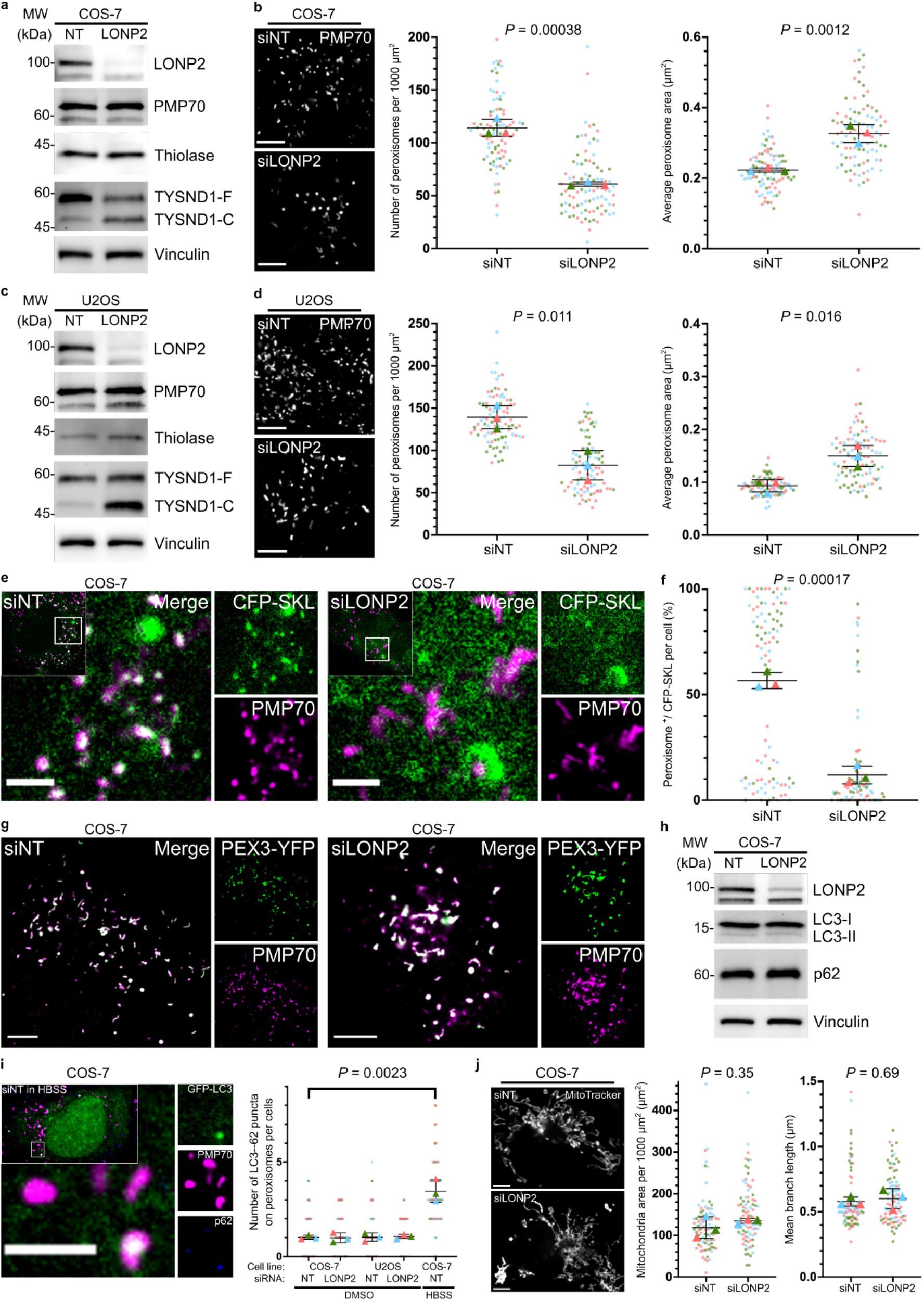
LONP2 knockdown triggered the accumulation of its substrate TYSND1, alterations in peroxisome size and numbers, and luminal protein import failure. **(a-d)** Immunoblotting (**a, c**) and representative confocal images and quantification of peroxisome size and numbers (**b, d**) in non-targeting (*NT*)- and *LONP2*-silenced COS-7 cells (**a, b**) and U2OS cells (**c, d**) after 144 h. TYSND1-F and TYSND1-C indicates full-length TYSND1 and the self-cleaved C-terminal region of TYSND1, respectively. Scale bar, 5 μm. (**e, f**) Representative confocal images (**e**) and quantification (**f**) of the colocalization between peroxisomes (PMP70) and transiently expressed peroxisome luminal protein (CFP-SKL) in *NT*- and *LONP2*-silenced COS-7 cells after 144 h. Scale bar, 2 μm. (**g)** Representative confocal images showing the colocalization between peroxisomes (PMP70) and transiently expressed peroxisome membrane protein (PEX3-YFP) in *NT*- and *LONP2*-silenced COS-7 cells after 144 h. Scale bar, 5 μm. (**h**) Immunoblotting for autophagy related proteins, vinculin was used as a loading control. (**i**) Representative confocal images of COS-7 cells grown in HBSS for 24 h (left) and quantification (right) showing the number of LC3-P62 puncta on peroxisomes of *NT*- and *LONP2*-silenced COS-7 cells and U2OS cells after 144 h. Cells were grown in either DMEM or HBSS for 24 h and LC3-P62 puncta on peroxisome was visualized with transiently expressed GFP-LC3 and the immunofluorescence of p62 and PMP70. Scale bar, 2 μm. (**j)** Representative confocal images and quantification showing the size and mean branch length of mitochondria in *NT*- and *LONP2*-silenced COS-7 cells after 144 h. Scale bar, 5 μm. On all plots, dots represent individual cells from *n* = 3 biologically independent experiments depicted in different colours, and triangles represent mean within an independent experiment. The mean values were used to calculate the average (horizontal bar), s.d. (error bars), and *P*-values (using two-tailed unpaired Student’s *t*-test).

The mechanisms of peroxisomal lumenal and membrane protein import are distinct (Farre et al., 2019). To monitor import efficiency of lumenal proteins, we transiently transfected cells with a peroxisome lumen-targeting sequence serine-lysine-leucine (SKL) fused with the Cyan Fluorescent Protein (CFP) and labelled peroxisomes with peroxisomal membrane protein PMP70. CFP-SKL was efficiently targeted to peroxisomes in control cells, but no longer imported into peroxisomes in *LONP2*-silenced COS-7 cells (**Fig. 1e and f**) or U2OS cells (**Fig. S1**), demonstrating that the knockdown of LONP2 impaired peroxisomal luminal protein import. To evaluate the efficiency of peroxisomal membrane protein import, we used adenoviral expression of peroxisomal biogenesis factor 3 (PEX3) tagged with Yellow Fluorescent Protein (PEX3-YFP), whose import was unaltered in *LONP2*-silenced COS-7 cells (**Fig. 1g**). These data establish that depletion of LONP2 in both cell lines leads to alterations in proteolysis of substrates like TYSND1 and inhibits protein import of our reporter CFP-SKL into the peroxisomal lumen without affecting membrane protein insertion.

Loss of mitochondrial LONP1 leads to accumulation of aggregated matrix content and eventual initiation of autophagy-mediated organellar turnover, or mitophagy (Zurita Rendon and Shoubridge, 2018). Thus, we evaluated whether LONP2 leads to induction of pexophagy by monitoring levels and localisation of LC3, a marker of autophagic membranes, and autophagic cargo adapter p62. As a positive control, we induced non-selective autophagy, using HBSS media, starving the cells of essential amino acids (Sargent et al., 2016). This resulted in an increased number of LC3-p62 puncta on peroxisome (**Fig. 1i, left**). However, the number of peroxisomes colocalized with transiently expressed GFP-LC3 and the endogenous p62 was unchanged between non-targeted (*NT*-) and *LONP2*-silenced cells after 6 days (**Fig. 1i, right**). Moreover, LONP2 knockdown did not induce global autophagy, as LC3 isoforms and p62 remained unchanged in COS-7 cells (**Fig. 1h**). These data are consistent with the analysis of peroxisome morphology, which shows enlarged organelles, and immunoblots that showed no change in peroxisomal proteins like PMP70 and thiolase (**Fig. 1a-d**).

Peroxisomal import failure can lead to the targeting of peroxisomal proteins to mitochondria with detrimental consequences on mitochondrial function, metabolism, and cristae architecture (Nuebel et al., 2021). Specifically, abrogation of *Pex3* or *Pex19*, core components of peroxisomal membrane insertion, resulted in the mitochondrial import of a series of peroxisomal membrane, and some luminal, proteins (Nuebel et al., 2021). Although Pex3-YFP was correctly targeted to peroxisomes in cells lacking LONP2 (**Fig. 1g**), we further examined whether LONP2 knockdown altered mitochondrial morphology by using Mitochondrial Network Analysis (MiNa), a semi-automated ImageJ plugin (Valente et al., 2017). Confocal imaging of mitochondria stained with MitoTracker Deep Red FM demonstrated that mitochondrial network morphology remained unchanged upon silencing of LONP2 (**Fig. 1j**). Taken together, the loss of LONP2 leads to peroxisomal proteotoxic stress, and alters peroxisomal morphology and numbers without initiation of pexophagy or altering mitochondrial network morphology. Thus, this allows us to uncouple consequences of peroxisomal proteotoxic stress from secondary mitochondrial defects.

### Peroxisomal proteotoxic stress leads to ISR and ribosome biogenesis activation in a cell specific manner

To establish the overall cell responses to peroxisomal proteotoxic stress, we analysed transcriptomes of NT- and LONP2-silenced cells using RNA sequencing (RNAseq) in COS-7 and U2OS cells. Surprisingly, the gene expression response resulting from loss of LONP2 between the two lines were drastically different (**Fig. 2a and b, Table S1**), despite similar peroxisomal morphology and import phenotypes **(Fig. 1 c-g)**. Unsupervised principal component analysis showed that *NT*- and *LONP2*-silenced samples separated clearly in COS-7 but not U2OS line (**Fig. 2a**). In COS-7, expression levels of 3835 genes were downregulated and of 3215 genes were upregulated (|log2(FC)|>0.5, p.adj<0.05), while in U2OS expression levels of only 182 and 77 genes were down- and up-regulated, respectively (|log2(FC)|>0.5, p-value <0.01) (**Fig. 2b, Table S1**).

**Figure 2.**
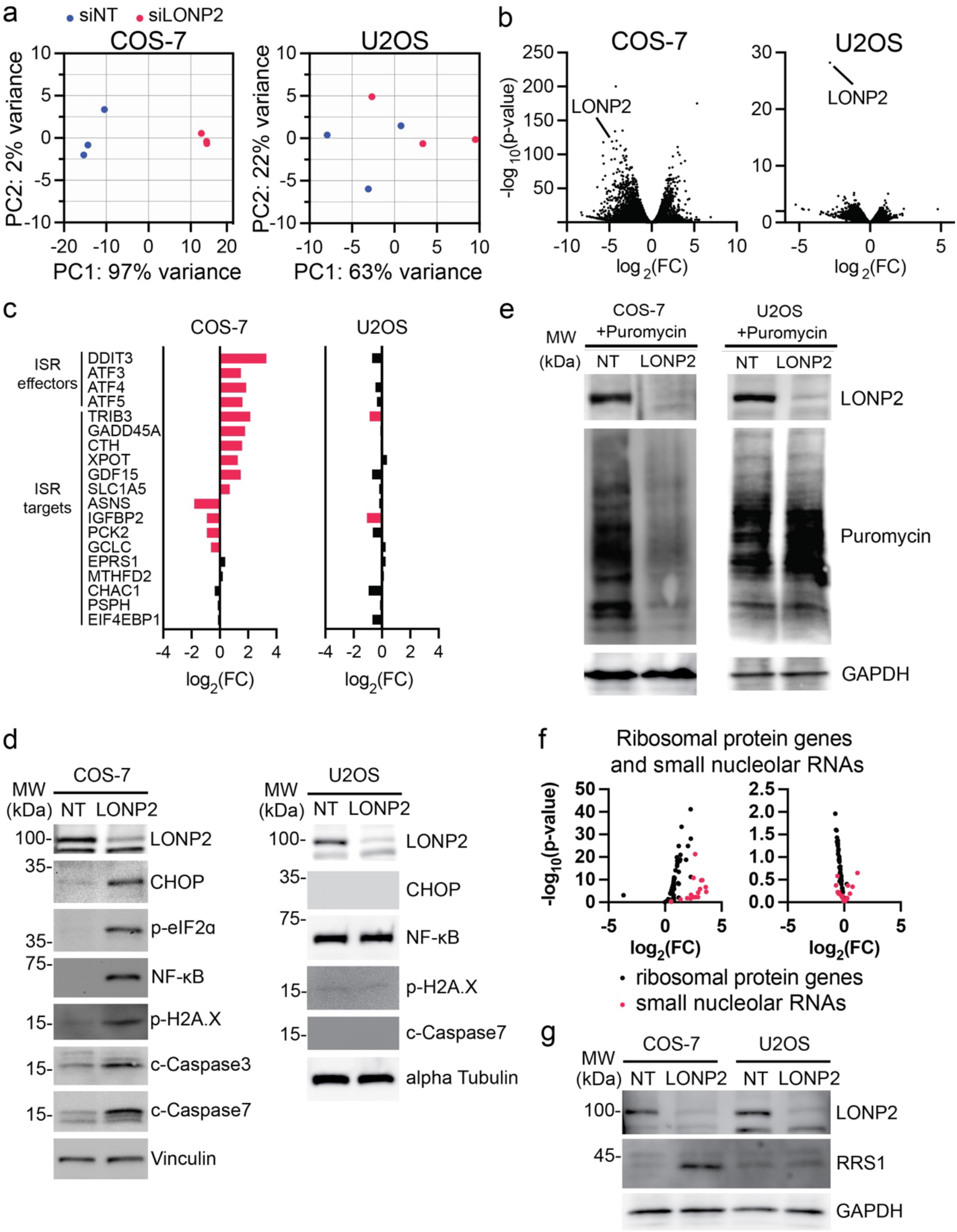
Cell-specific ISR activation upon silencing of LONP2. RNA sequencing (*n* = 3 independent experiments per condition), LONP2 silencing for 7 days. **(a)** Principal component (PC) score plot. **(b)** Volcano plots showing all identified transcripts. **(c)** Transcripts encoding integrated stress response (ISR) factors. **(d)** Immunoblotting for ISR related proteins, vinculin was used as a loading control. **(e)** Translation assay: cells were incubated with puromycin for 10 min., and incorporation of puromycin into newly synthetized proteins was revealed with anti-puromycin antibodies. GAPDH was used as a loading control. **(f)** Transcripts encoding ribosomal biogenesis related rRNA, snoRNA and mRNA genes. (**g**) Immunoblot of ribosomal biogenesis regulator RRS1 in *NT*- and *LONP2*-silenced COS-7 cells and U2OS cells.

As we anticipated a stress response signature upon loss of LONP2, we focused our analysis on the integrated stress response (ISR). The ISR is an evolutionary conserved eukaryotic signaling pathway that acts to attenuate cap-dependent translation via phosphorylation of eukaryotic translation initiation factor (eIF2), while simultaneously inducing one or more ISR effectors [activating transcription factors (ATF) ATF3, 4, 5, C/EBP homologous protein (CHOP) encoded by *DDIT3*] and their downstream targets (Costa-Mattioli and Walter, 2020; Pakos-Zebrucka et al., 2016). ISR effectors ATF3-5, CHOP, and several of their targets were induced in LONP2 depleted COS-7 but not in U2OS cells (**Fig. 2c and d**). Considering that one of the hallmarks of ISR is downregulation of mRNA translation we examined the effects of LONP2 depletion on protein synthesis by labeling cells with puromycin for 10 minutes (Schmidt et al., 2009). These experiments revealed a dramatic reduction in protein synthesis in LONP2-depleted as compared to control COS-7 cells (**Fig. 2e**). In contrast, LONP2-depletion failed to induce CHOP levels and attenuate protein synthesis in U2OS cells, also consistently with the RNA signature (**Fig. 2d and e**). Induction of global cellular stress in LONP-2 depleted COS-7 cells was further illustrated by the activation of apoptosis-associated cleaved caspases 3 and 7, increased DNA damage, as monitored by γ2HAX phosphorylation and induction of a regulator of innate immunity transcription factor NF-kB (**Fig. 2d**). These effects were not observed upon LONP2 depletion in U2OS cells. Together, these data suggest that proteotoxic peroxisomal stress activates ISR in a cell-type specific manner.

We also observed a global upregulation of mRNAs encoding factors implicated in ribosome biogenesis in LONP2-depleted COS-7, but not U2OS cells (**Fig 2f**). Expression of a core mediator of ribosomal assembly RRS1 was also increased at the protein level **(Fig. 2g)** (Hua et al., 2021). This was unexpected, since ribosome biogenesis is generally suppressed under stress (Lindstrom et al., 2022; Ni and Buszczak, 2023; Szaflarski et al., 2022), and LONP2-depletion in COS-7 cells resulted in strong reduction in protein synthesis (**Fig. 2e**). Collectively, these findings suggest that in addition to induction of ISR, depletion of peroxisomal LONP2 results in altered ribosome biogenesis in COS-7 cells.

A core goal of stress responses to organellar dysfunction is to restore homeostasis, which often includes an upregulation of chaperone proteins, and in this case, potentially peroxisomal proteins. A previous report in *C*.*elegans* characterizing the response to PEX5 deletion noted an upregulation of peroxisomal genes driven by the activation of transcription factors PPARα/MED15 orthologues (Rackles et al., 2021). However, in both COS-7 and U2OS, depletion of LONP2 did not result in an upregulation of peroxins or peroxisomal genes (**Fig. S2**).

### Peroxisomal proteotoxic stress leads to defective retinoic acid signalling

While the ISR was not induced in U2OS cells, we sought to identify common responses to peroxisomal proteotoxic stress between COS-7 and U2OS cells. Gene ontology analysis of transcripts changed in both cell lines revealed a significant downregulation of the retinoic acid (RA) / vitamin A signalling pathway (Petkovich and Chambon, 2022) as a top hit (**Fig. 3a**). We validated this at the protein level by immunoblotting key proteins involved in RA signaling pathway: cellular all-trans retinoic acid (ATRA) binding proteins (CRABP2) and aldehyde dehydrogenase 1 family member A2 (ALDH1A2) (Pohl and Tomlinson, 2020) (**Fig. 3b**). Consistent with the RNAseq data, CRABP2 levels were strongly reduced in both COS-7 and U2OS. However, a second enzyme on this pathway ALDH1A2 was not changed in U2OS, suggesting that the effects at the protein level in this cell line may be partial.

**Figure 3.**
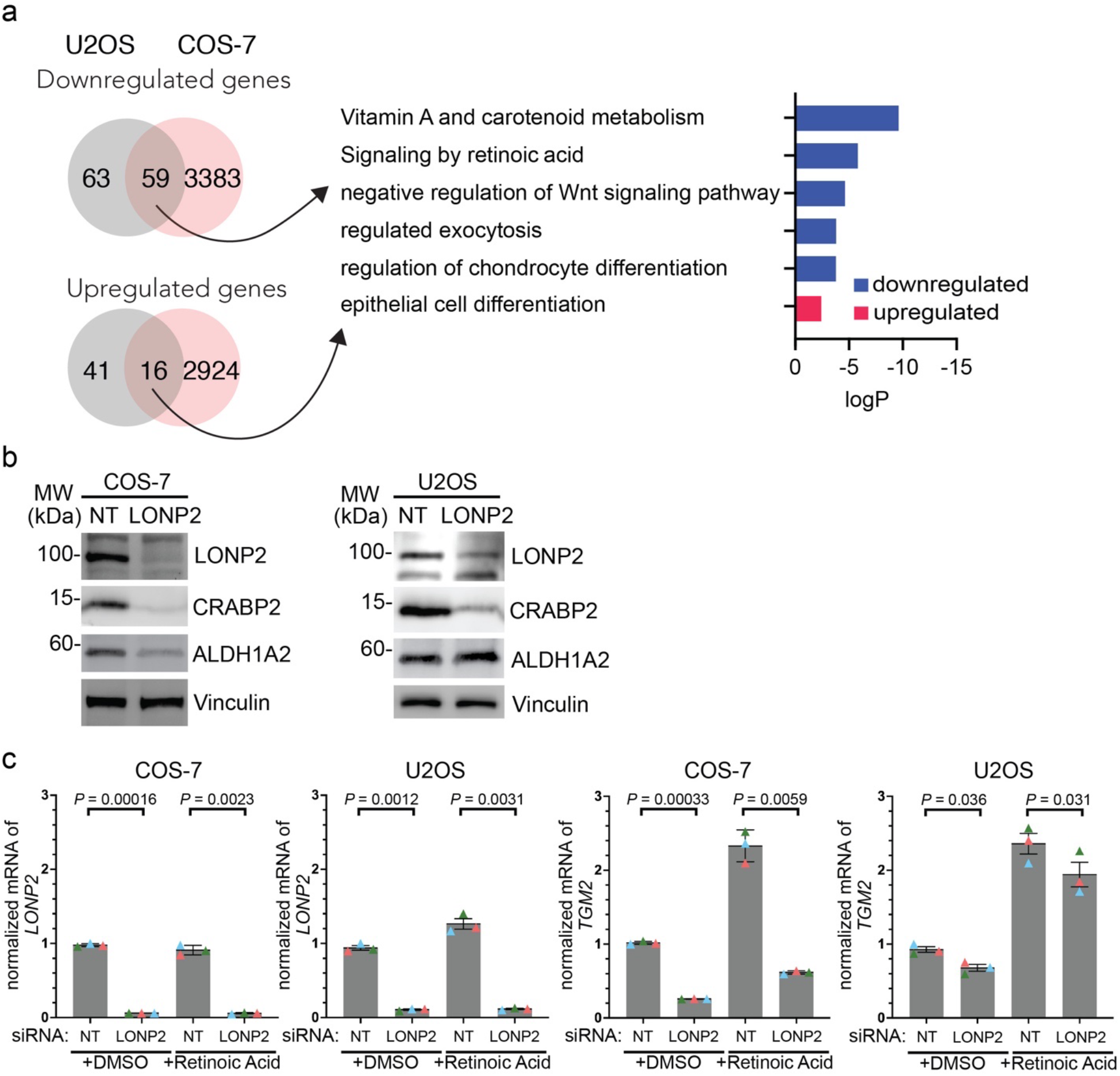
LONP2 silencing leads to impaired retinoic acid signaling. **(a)** Gene ontology of RNA sequencing data (dataset is described in Fig. 2). Numbers on Venn diagram indicate numbers of genes in each category. (**b**) Immunoblotting of retinoic acid regulating proteins, vinculin used as a loading control. (**c**) COS-7 or U2OS cells were incubated in presence or absence of 5 μM retinoic acid for 24 h and mRNA transcript levels for LONP2 (left two panels) and TGM2 (right two panels) quantified with qRT-PCR. *n* = 3 independent experiments, means are depicted in different colours. The mean values were used to calculate the average (horizontal bar), s.d. (error bars) and *P*-values (using two-tailed unpaired Student’s *t*-test).

To test the functional consequence of the downregulation of RA signaling enzymes, we treated cells with all-*trans* retinoic acid (ATRA) to activate RA signaling and drive transcription of specific gene targets. RT-qPCR analysis confirmed the efficiency of LONP2 silencing in both cell lines (**Fig. 3c**). Upon ATRA treatment, the transcriptional upregulation of a retinoic acid-responsive gene, *Transglutaminase 2* (*TGM2*), was strongly suppressed in LONP2 knockdown in COS-7 cells, but only mildly altered in U2OS (**Fig. 3c**). Indeed, levels of TGM2 mRNA were reduced in COS-7 cells treated with siLONP2 even prior to ATRA stimulation. These data demonstrate the impact of peroxisomal dysfunction on retinoic acid signaling pathways.

### Lipidomic analysis revealed accumulation of sphingomyelins and cholesterol esters in LONP2 silenced cells

One of the core functions of peroxisomes is in the generation and catabolism of specific lipids. Therefore, to investigate the effect of LONP2 knockdown on the regulation of lipid homeostasis, we performed comprehensive untargeted lipidomics in both COS-7 and U2OS cells. The dataset retained 2,094 MS features, defined by their m/z, RT and signal intensity. Similar to gene expression data, the lipidomic changes resulting from loss of LONP2 between the two lines were drastically different as can be seen from the score plot of unsupervised principal component analysis (**Fig. 4a**). Among the 2,094 features of the final dataset, 467 (22%) of them passed our selected threshold (|(FC)| > 1.5, *P*_*corr*_-value < 0.05) as significantly different between the two cell lines under basal condition (**Fig. S3a, Table S2**), whereas upon depletion of LONP2 (vs. siNT), 242 were changed in COS-7 and 234 in U2OS cells (**Fig. S3b**). It is notable that the lipid profile differed more markedly between the two cell lines at baseline (∼22% were significantly different) than upon silencing of LONP2 within in each cell line (∼11% significantly different).

**Figure 4.**
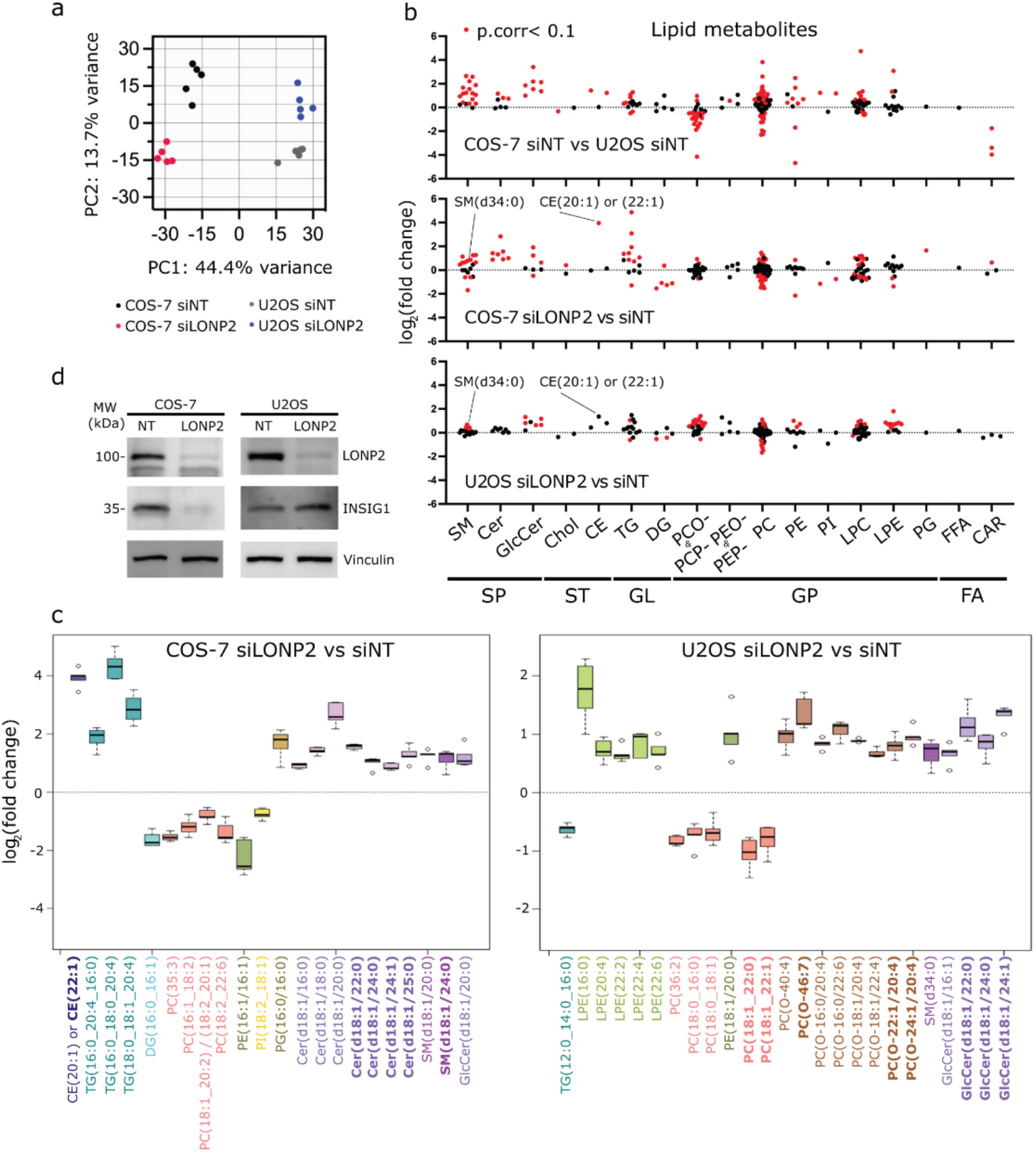
Lipid profiles of COS-7 and U2OS cells differ drastically and are differentially affected by LONP2 silencing. Untargeted lipidomic analysis of *n* = 5 independent cell experiments for each group under basal condition of after *LONP2*-silencing for which the final dataset included 2,094 lipid features (Table S2). **(a)** Principal component (PC) score plot. **(b)** Log_2_ (fold changes) of 206 lipid features annotated by MS/MS and data alignment using our in-house database, and grouped by lipid subclass, whereby red and black dots indicate lipids that reach or not our selected significance threshold for the indicated comparison, *P*_*corr*_-value < 0.1. (**c)** Box plots of unique lipids annotated by MS/MS from the 100 most significant lipid features discriminating *LONP2-*silenced COS-7 cells (top) and U2OS cells (bottom) from *NT-*silenced cells. The midline represents the median fold change vs. *NT-*silenced cells, the box represents the interquartile range between the first and third quartile, and whiskers represent the lowest or highest values. (**d**) Immunoblotting of cholesterol sensor INSIG1, vinculin used as a loading control. Abbreviations: sphingolipid (SP), sterol lipid (ST), glycerolipid (GL), glycerophospholipid (GP), fatty acyls (FA), sphingomyelin (SM), ceramide (Cer), glucosylceramide (GlcCer), cholesterol (Chol), cholesterol ester (CE), triacylglycerol (TG), diacylglycerol (DG), 1-alkyl, 2-acylglycerophosphocholine (PCO-), 1-(1Z-alkenyl), 2-acylglycerophosphocholine (PCP-), 1-alkyl, 2-acylglycerophosphoethanolamine (PEO-), 1-(1Z-alkenyl), 2-acylglycerophosphoethanolamine (PEP-), diacylglycerophosphocholine (PC), diacylglycerophosphoethanolamine (PE), diacylglycerophosphoinositol (PI), monoacylglycerophosphocholine (LPC), monoacylglycerophosphoethanolamine (LPE), diacylglycerophosphoglycerol (PG), free fatty acid (FFA) and acylcarnitine (CAR). All lipid features were categorized into subclasses and labeled based on the LIPID MAPS^®^ Structure Database.

Given the untargeted nature of lipidomic analysis, only lipid features of interest have been annotated, which for this study are those that passed the selected threshold of significance for the impact of LONP2 silencing in COS-7 and U2OS cell lines (See Methods for details). A total of 206 lipid features were annotated using MS/MS and data alignment with our in-house database (Forest et al., 2018) and were found to belong to different lipid subclasses (**Fig 4b**). The most significantly changed unique lipid species upon LONP2 silencing in COS-7 cells and U2OS cells, which were annotated using MS/MS (22 and 25 unique lipids, respectively), are shown as box plots (**Fig 4c**).

From these data, we conclude that for all three group comparisons conducted in this study, glycerophospholipids, specifically phosphatidylcholines (PC), showed the highest number of significantly up- or downregulated lipid features (red dots; Figure 4b), reflecting substantial remodeling of their fatty acyl side chains. Under the basal state, sphingolipids, both sphingomyelins and glucosylceramides, as well as cholesterol esters were more abundant in COS-7 cells compared to U2OS cells (SP and ST; Fig. 4B). Upon LONP2 silencing, COS-7 and U2OS cells shared some lipid changes: namely downregulation of PCs (**Fig. 4b and orange boxes in Fig. 4c**), upregulation of sphingolipids, namely sphingomyelins (SM) such as SM(d34:0) and glucosylceramides (**SP in Fig. 4b; Purple boxes in Fig. 4c; Fig. S3c**) and of lipids containing a saturated or monounsaturated very long chain fatty acyl side chain (VLCFA ≥22 carbon chains in bold, **Fig. 4c**).

There were, however, important differences between these two cell lines in the response to LONP2 silencing for other lipid subclasses. For example, and of relevance to cholesterol metabolism, cholesterol ester (CE) CE20:1 or CE22:1 showed a striking ∼4 log_2_(FC) accumulation in *LONP2*-silenced COS-7 cells (*P*_*corr*_ = 0.000091). The accumulation of these CEs in *LONP2*-silenced U2OS cells was less prominent, and variable (upregulated in 2 samples, unchanged in 3), therefore did not pass our significance threshold (log_2_(FC)=1.36, *P*_*corr*_ = 0.26) (**Figure 4b and c, Fig. S3c; and Table S2**). A prominent set of ceramide species were also upregulated in COS7 cells, many of which carried acyl chains ≥22 carbons. In contrast, ether PC were upregulated in *LONP2*-silenced U2OS but not COS-7 cells (**Brown boxes in Fig. 4c**).

Given the striking accumulation of CEs in *LONP2*-silenced COS-7 cells vs. *LONP2*-silenced U2OS cells, we further investigated the sensing of the excess cholesterol in our system. For this, we analysed expression of the ER membrane protein insulin induced gene 1 (INSIG1), a core sensor within the cholesterol biosynthetic pathway that is rapidly degraded in response to cholesterol depletion (Brown et al., 2018; Radhakrishnan et al., 2008). Surprisingly, INSIG1 showed downregulation with immunoblotting (**Fig. 4d**) and RNAseq (log_2_(FC) = 3.34, p.adj = 2.56E-14) in COS-7 cells. In U2OS, INSIG1 mRNA and protein remained unchanged (**Fig. 4d**; RNAseq log_2_(FC) = 0.17, p.value = 0.44). Thus, concurring with lipidomic data, these results show that in COS-7 cells, cholesterol is accumulated upon peroxisomal proteotoxic stress, but the loss of INSIG1 suggests that ER cholesterol sensing is impaired.

### Cholesterol flux analysis shows accumulation in endolysosomes in response to LONP2 depletion

Impaired sensing of excess cholesterol in LONP2 depleted cells prompted us to examine the subcellular localisation of the elevated cholesterol. For this, we stained cholesterol with Filipin. Confocal imaging showed cholesterol staining to be abundant in both Nile Red-stained lipid droplets, and endolysosomes marked with lysosomal-associated membrane protein 1 (LAMP1) (**Fig. 5a-b**). LONP2 knockdown also led to increased number of lipid droplets (**Fig. 5a**), consistent with increased triglycerides seen in lipidomics analysis (**TG in Fig. 4b**).

**Figure 5.**
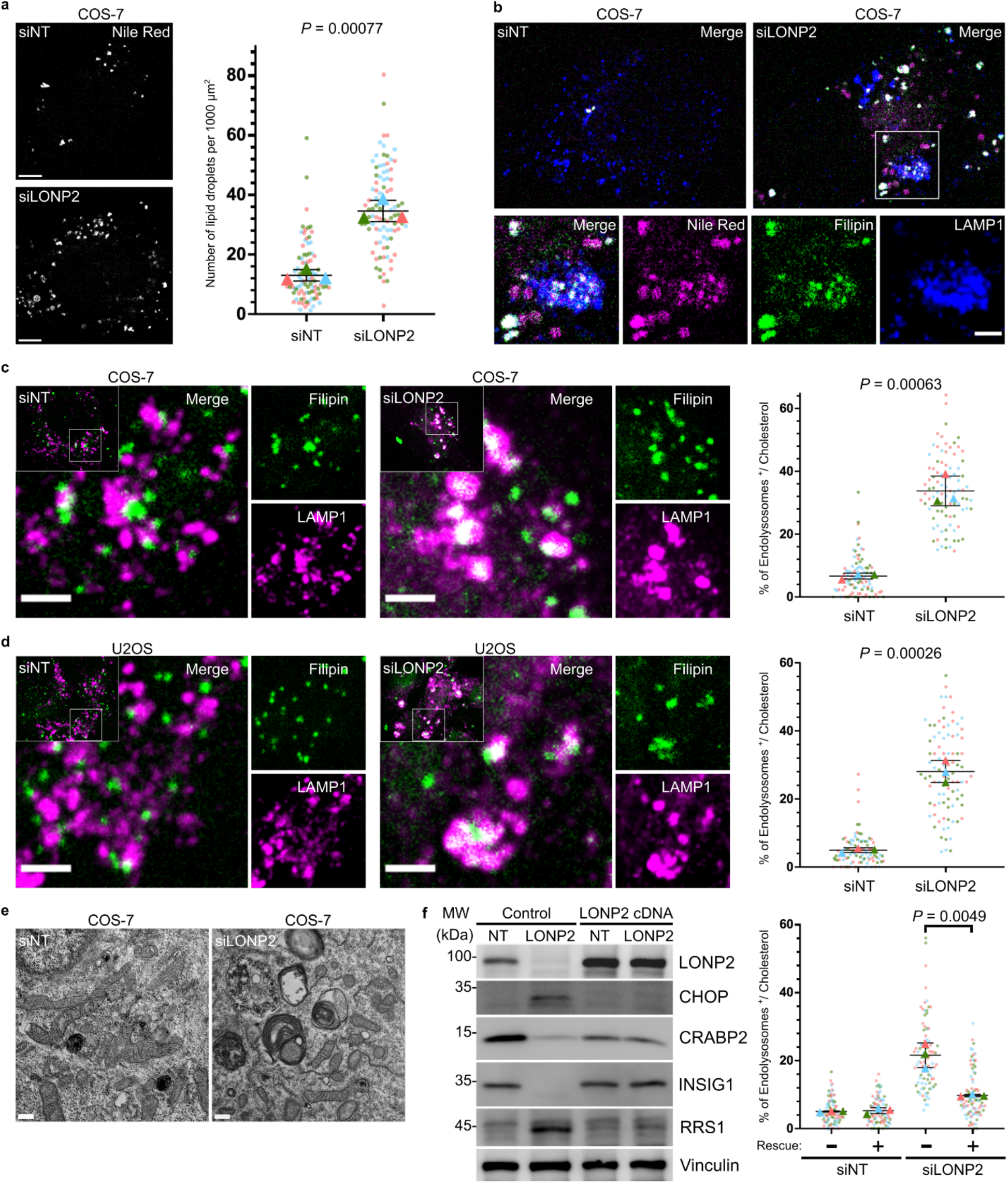
LONP2 knockdown induced lipid storage, including aberrant cholesterol accumulation in lysosomes. **(a)** Representative confocal images (left) and quantification (right) of lipid droplets (Nile Red) in *NT*- and *LONP2*-silenced COS-7 cells after 6 days. Scale bar, 5 μm. **(b)** Representative confocal images showing the Filipin-stained intracellular cholesterol (green), Nile Red-stained lipid droplets (magenta) and endolysosomes (blue) visualized with the immunofluorescence of LAMP1. Scale bar, 2 μm. (**c, d)** Free cholesterol overload experiment in *NT*- and *LONP2*-silenced COS-7 cells (**c**) and U2OS cells (**d**) after 7 days. Representative confocal images (left) and quantifications (right) of the colocalization between cholesterol (Filipin) and endolysosomes (LAMP1). Scale bar, 2 μm. (**e)** Transmission electron microscopy micrographs of *NT*- and *LONP2*-silenced COS-7 cells. Scale bar, 500 nm. **(f)** Immunoblotting (left) and image quantification (right) of cholesterol overload experiment in *NT*- and *LONP2*-silenced COS-7 cells rescued with transient expression of siRNA-resistant LONP2 cDNA for 48 h. Data from *n* = 3 biologically independent experiments, and their respective means are depicted in different colours. The mean values were used to calculate the average (horizontal bar), s.d. (error bars) and *P*-values (using two-tailed unpaired Student’s *t*-test).

To monitor the flux of cholesterol in the absence of LONP2, we incubated cells with excess free cholesterol for one hour and stained with Filipin. As with the endogenous cholesterol, confocal microscopy showed enhanced accumulation of Filipin within endolysosomes in both *LONP2*-silenced COS-7 and U2OS cells as compared to corresponding controls (**Fig. 5c and d, left**). Filipin staining was observed in less than 8% of endolysosomes in control COS-7 or U2OS cells, which rose to approximately 30% of endolysosomes in *LONP2*-silenced COS-7 and U2OS cells (**Fig. 5 c and d, right**). The accumulation of cholesterol in lysosomes is reminiscent of the phenotypes in human lysosomal storage diseases, such as Niemann-Pick disease type C (NPC) caused by pathogenic variants in *NPC1* or *NPC2*, which block cholesterol export from the lysosome (Pfeffer, 2019). Indeed, electron microscopic analysis of *LONP2*-silenced COS-7 cells revealed many lamellated structures and electron-dense particles that strongly resembled the abnormal lysosomes observed in NPC cells (**Fig. 5e**). This included the presence of lipid whorls and lamellar inclusions, consistent with observation that LONP2 depletion impaired sphingomyelin and cholesterol homeostasis (**Fig. 5e**).

Our results indicate that LONP2 knockdown impairs the ability of cholesterol to efflux from endolysosomes. Trapping of cholesterol in endolysosomes may lead to cholesterol depletion within the ER, driving INSIG1 degradation (**Fig. 4d**). This was specific to COS-7 cells, which started with significantly higher levels of sphingomyelins, glucocylceramides and cholesterol esters relative to U2OS cells. This may indicate that the steady state flux of cholesterol and sphingomyelins are much higher in COS-7 at steady state, rendering them more sensitive to LONP2 depletion. Indeed, only upon addition of excess cholesterol for flux analysis in U2OS cells did we observe similar phenotypes to COS-7 cells with the accumulation of cholesterol in endolysosomes. This is consistent with previous work suggesting that peroxisomes are required for the flux of cholesterol from the late endosome to the ER (Chu et al., 2015). Taken together, our data establish a critical requirement for functional peroxisomes in the trafficking of cholesterol and lipid droplet biogenesis. Interestingly the cholesterol trafficking defect was observed in both U2OS and COS-7 cells, which indicates a phenotype independent of the ISR since LONP2 depletion induced ISR only in COS-7 but not U2OS cells.

To exclude the possibility of off-target effects in our LONP2 silencing experiments, COS-7 cells were transfected with different siRNA targeting the 3’ UTR of *LONP2* and then rescued by transient expression of si-resistant LONP2 cDNA. Immunoblotting revealed that CHOP activation, as well as downregulation of INSIG1 and CRABP2 were fully rescued by the transient expression of LONP2 cDNA (Fig. 5f, left). Cholesterol homeostasis was also rescued functionally, as free cholesterol no longer accumulated at endolysosomes after transfection of LONP2 cDNA for 48 h (Fig. 5f, right).

### Specificity of response to siLONP2 relative to loss of Pex5

Previous approaches to investigate peroxisomal stress pathways focused on loss of core peroxins essential for protein import in *C. elegans*, Drosophila and mammalian cells (Mast et al., 2011; Nuebel et al., 2021; Rackles et al., 2021; Wangler et al., 2017). LONP2 is a chaperone and protease that is important for the turnover and proteostasis within peroxisomes, and, as we demonstrated here, is required for the import of at least some luminal proteins. Therefore we tested whether the pathways that are induced by LONP2 silencing are also triggered by the loss of core import proteins like PEX5 in our system. As with the loss of LONP2, PEX5-depleted COS-7, but not U2OS cells displayed clear upregulation of ISR effector CHOP, as well as downregulation of the ER cholesterol sensor INSIG1 (**Fig. S4**). In contrast to the depletion of LONP2 however, PEX5 silencing failed to bolster expression of the ribosomal assembly protein RRS1 or retinoic acid CRABP2 protein levels (**Fig. S4**). These intriguing differences suggest that luminal peroxisomal proteotoxic stress induces distinct responses relative to global peroxisomal protein import failure.

## Discussion

Data presented here extend and identify new consequences of peroxisomal dysfunction, including induction of the ISR, disruption of cholesterol regulation and trafficking, and altered retinoic acid signalling. Our data highlight the significant variability in the dependence of different cell types on peroxisomal function. Importantly, our strategy to manipulate the peroxisomal protease/chaperone LONP2 allowed us to maintain peroxisomal membrane protein import and thus study peroxisomal stress response without the induction of mitochondrial stress (Nuebel et al., 2021).

Our data showed that loss of LONP2 led to changes in peroxisome morphology, where cells had fewer and larger organelles, suggesting that LONP2 function was directly or indirectly required for peroxisomal division. While we expected to observe an induction of pexophagy as part of a peroxisomal quality control response, we saw no induction of autophagy or changes in peroxisomal mass. This may be explained, at least in part, by the global alterations in lysosomal lipid accumulation shown in previous work to interfere with autophagy (Sarkar et al., 2013). It is also plausible that LONP2 function is itself required for the induction of pexophagy, or perhaps pexophagy is not the major pathway in peroxisomal quality control. We were also surprised not to see a transcriptional induction of peroxisomal biogenesis, a repair mechanism seen in previous investigations into the peroxisome stress response pathways (Rackles et al., 2021).

Lipidomics, RNAseq and cell biological analysis presented here further establish peroxisomes as a critical contributor to cholesterol homeostasis and intracellular flux. This is consistent with recently published evidence that peroxisomes form direct contact sites with lysosomes to flux cholesterol to the ER (Chu et al., 2015). To this end, synaptotagmin7 (syt7) was identified as the essential tether between peroxisomes and lysosomes to drive cholesterol flux. We observed no apparent changes in syt7 mRNA levels (**Table S1**). We thus suspect that the loss/aggregation of an unidentified peroxisomal luminal protein interfered with this pathway of cholesterol flux. The loss of INSIG1 was also seen upon loss of PEX5 in COS-7 cells, indicating that impaired cholesterol trafficking may occur upon diverse peroxisomal dysfunctions. Untargeted lipidomics also revealed an increase in mono- or unsaturated very long chain fatty acyl chains among the significantly changed lipids, suggesting peroxisomal dysfunction in beta-oxidation pathways upon depletion of LONP2. Interestingly, these changes were enriched within ceramide and sphingomyelin species in COS7, but within ether lipids and glucocylceramides in U2OS. This is further indication that peroxisomes play important, yet differential roles in the wiring of lipid metabolism between the two cell lines. The impact of these elevated lipids on cell survival may be reflected in the increased caspase activation in COS7, where ceramides are tightly linked to enhanced apoptosis (Nganga et al., 2018). The increased ether lipids may alter susceptibility of U2OS cells to ferroptotic death pathways, as highlighted in recent studies linking peroxisomal function to ferroptosis (Zou et al., 2020). Future work will continue to explore the contribution of peroxisomal lipid metabolism to cell survival pathways.

Another common pathway altered upon loss of LONP2 (but not PEX5) in both cell lines was a prominent downregulation of the retinoic acid/Vitamin A pathway. Retinoic acid is a branched chain lipid that plays a key role in a plethora of both essential in development, and disease-related pathways (Pohl and Tomlinson, 2020). Historically, retinoic acid has been described as a peroxisome proliferator (Hertz and Bar-Tana, 1992) and a possible link between peroxisomes and lipid droplets (Robison and Kuwabara, 1977). Intriguingly, RA signaling pathway has been linked to peroxisomal function in liver, specifically to bile acid and cholesterol metabolism (Saeed et al., 2017). Retinoic acid binds to the nuclear receptor RXR, which controls gene expression of a host of genes and can form dimers with up to 10 other nuclear hormone receptors including regulators of bile acid metabolism and lipid homeostasis (Petkovich and Chambon, 2022). Of relevance to peroxisomes, knockout of RXRα in mouse hepatocytes leads to a 7 fold upregulation of a key regulator of bile acid metabolism CYP7A1, again linking retinoic acid signaling and peroxisomal function (Wan et al., 2000). Our data suggests a retrograde signaling pathway from peroxisomes into the RXR signaling pathway as well. Future work will explore the physiological, tissue specific regulation to unravel how and why the loss of LONP2 results in the downregulation RA signaling.

Additional responses to LONP2 depletion included the COS-7-specific induction of the ISR, and the increased mRNA transcripts linked to ribosome biogenesis. The ISR is a highly conserved stress response pathway, classical triggers of which include amino acid or heme deprivation, viral infection, and ER stress (Pakos-Zebrucka et al., 2016). ISR is seen as a pro-survival mechanism, however recent studies demonstrated the context-specific and time-sensitive role of the response in a variety of pathologies including models of neurodegenerative and mitochondrial diseases (Costa-Mattioli and Walter, 2020; Kaspar et al., 2021; Spaulding et al., 2021). Our study establishes peroxisomal proteotoxic stress as a new ISR trigger, providing a potential target for therapeutic interventions. COS-7 cells lacking LONP2 also demonstrated increased transcription of ribosome biogenesis messages that included rRNA, snoRNA and mRNAs. Interestingly, PEX5 loss did not lead to the increased expression of RRS1, a core regulator of ribosomal biogenesis, yet showed a clear activation of the ISR with CHOP expression. Another recent study showed significant downregulation of mRNA linked to ribosome biogenesis upon loss of peroxins (Huang et al., 2021), a trend also seen in our Pex5 depletion experiment. Therefore, the ribosome biogenesis response can be uncoupled from the peroxisome-driven activation of the ISR and appears to be specific to loss of LONP2.

The acute silencing approach allowed us to observe the immediate response to the loss of a core peroxisomal chaperone/protease, avoiding the confounding effects of cellular adaptation in complete loss of function models. Our data suggest that the cell-specific responses reflect a differential dependence on peroxisomal functions in lipid handling between cell types and tissues. Peroxisomes exhibit profound functional plasticity depending on the tissue context, driving bile acid synthesis in liver to the generation of ether lipids in oligodendrocytes, and scavenging ROS (Wanders et al., 2023). It will be important to further explore the consequences of peroxisomal stress in tissues and disease states, where the study of dynamic lipid handling and flux may be studied in more physiological settings. Understanding the contributions of peroxisomes to cholesterol flux in liver during bile acid synthesis, or within development/cancer model systems where peroxisomes may impact retinoic acid metabolism, could open doors to new therapeutic targets in multiple disease settings from Zellwegers syndrome to metabolic disorders and cancer.

## Acknowledgements

We are grateful for support from Sigrid Juselius Fellowship (OI), FRQS Studentship (AY) and a CRC (HMM). This work was funded by CIHR PJT-162183 (to HMM). We would like to thank members of the laboratory for critical reading of the manuscript.

## Author contributions

AY designed and performed experiments, contributed to manuscript preparation

OI contributed to analysis and presentation of both RNAseq and lipidomics datasets, helped design validation experiments, and contributed to the preparation of the manuscript.

MN performed experiments and assisted AY in this work

RL performed RNAseq experiments, and KW provided analysis of RNAseq dataset

CD, IRF and CDR performed lipidomics experiments, analyzed and interpreted the data

IT provided expertise for puromycin experiments and expert advice related to ISR, ribosome assembly and translation.

HMM designed and analyzed experiments, contributed to manuscript and figure preparation.

All members of the team contributed to manuscript preparation and editing.

## Supplementary Figure Legends

**Figure S1.**
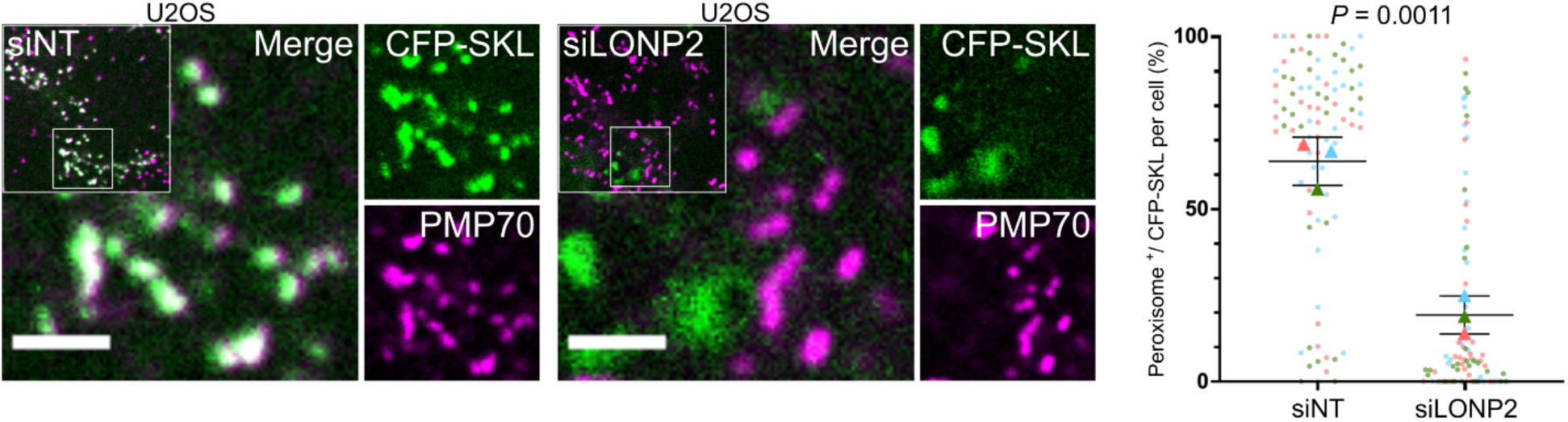
LONP2 knockdown triggered luminal protein import failure in U2OS cells. Representative confocal images (left) and quantification (right) of the colocalization between peroxisomes (PMP70) and transiently expressed peroxisome luminal protein (CFP-SKL) in non-targeting (NT)- and *LONP2*-silenced U2OS cells after 144 h. Scale bar, 2 μm. Dots represent individual cells from *n* = 3 biologically independent experiments depicted in different colours, and triangles represent their respective means. The mean values were used to calculate the average (horizontal bar), s.d. (error bars), and *P*-values (using two-tailed unpaired Student’s *t*-test).

**Figure S2.**
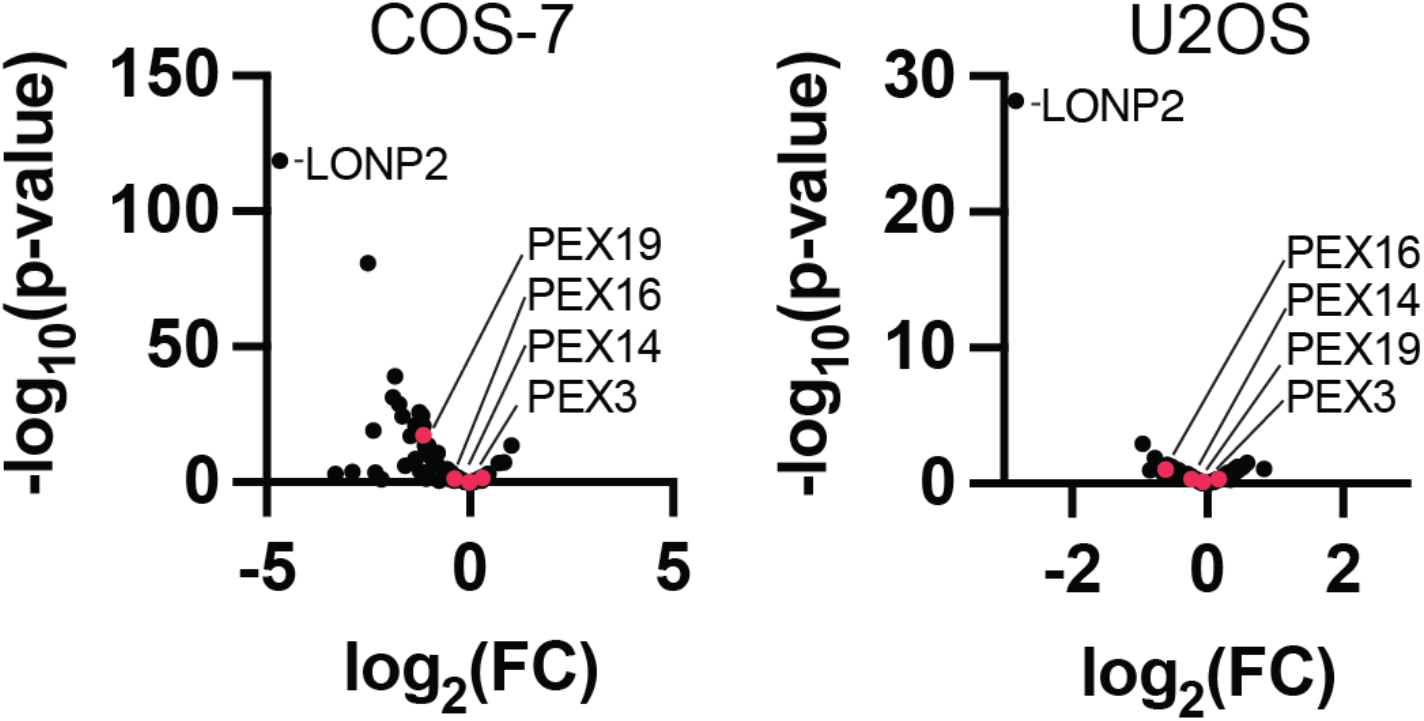
Peroxisomal gene expression is not significantly altered upon loss of LONP2. Volcano plots showing expression of peroxisomal genes in both COS-7 (left) and U2OS (right). Selected peroxins linked to peroxisomal biogenesis are highlighted with red dots.

**Figure S3.**
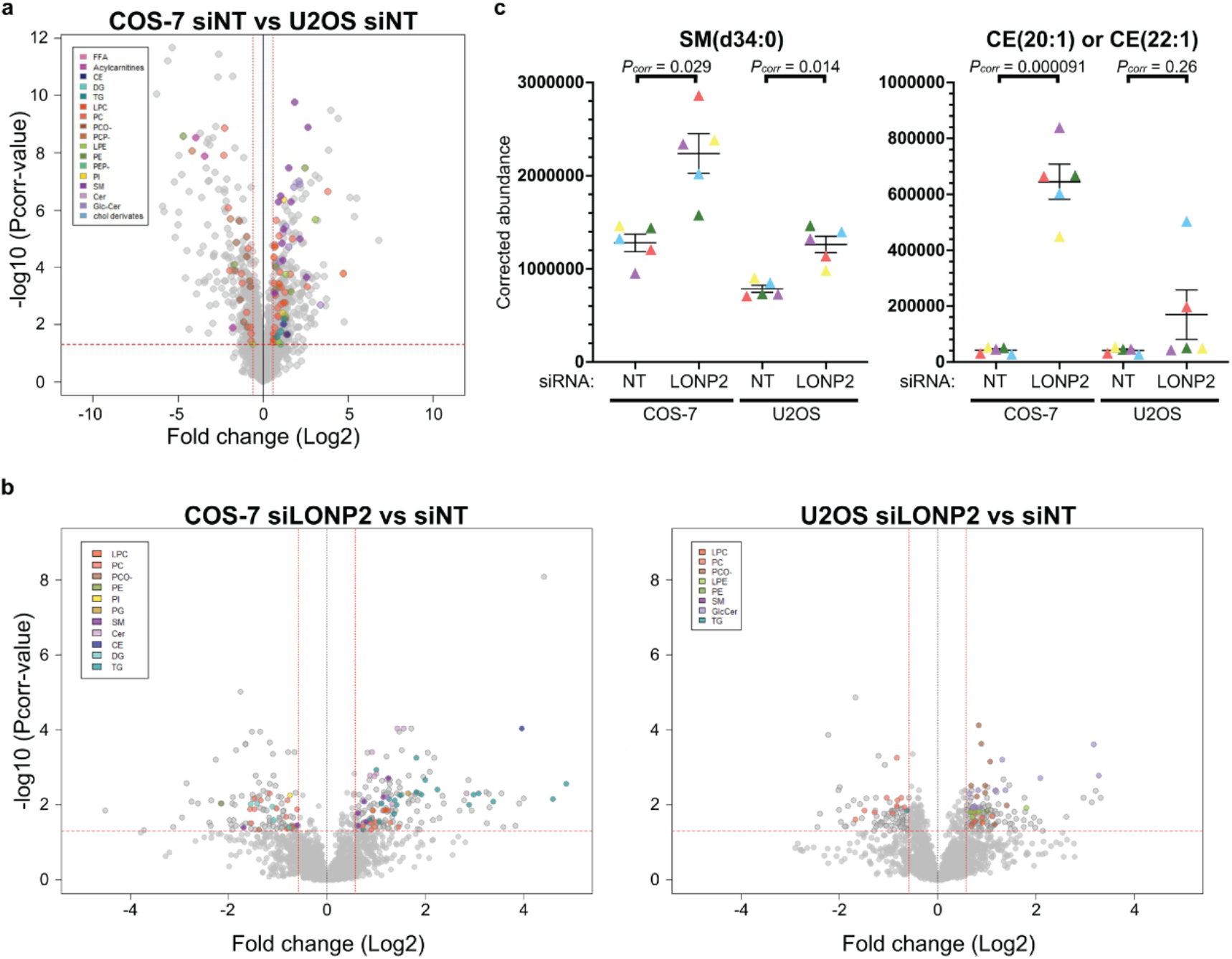
Differences and commonalities of lipidomic changes between COS-7 cells and U2OS cells. Volcano showing all 2,094 lipid features retained in the final dataset for the comparison between COS-7 and U2OS cells under steady state conditions (**a**) and 7-day of *LONP2*-silenced vs. NT-silenced COS-7 cells (**b, left**) or U2OS cells (**b, right**). X-axes show fold-change (FC) (log_2_) in MS signal intensity values and y-axes the corresponding corrected p-values (-log_10_*P*_*corr*_) following testing using unpaired Student *t*-test followed by a Benjamini-Hochberg correction. Lipid features that significantly passed our selected thresholds as indicated by red dotted lines (horizontal: *P*_*corr*_ value <0.05; and vertical |(FC)|> 1.5)) were annotated to unique lipids using MS/MS and data alignment with an in-house database (see Table S2 for details). Colors indicated lipid subclasses as indicated in the small inset. Grey dots indicate features that have not been identified. (**c)** Dot plots for sphingomyelin (SM(d34:0)) and cholesterol ester (CE(20:1) or CE(22:1)) in both COS-7 cells and U2OS cells. Data from *n* = 5 biologically independent experiments. Abbreviations: free fatty acid (FFA), diacylglycerol (DG), triacylglycerol (TG), monoacylglycerophosphocholine (LPC), diacylglycerophosphocholine (PC), 1-alkyl, 2-acylglycerophosphocholine (PCO-), 1-(1Z-alkenyl), 2-acylglycerophosphocholine (PCP-), monoacylglycerophosphoethanolamine (LPE), diacylglycerophosphoethanolamine (PE), 1-(1Z-alkenyl), 2-acylglycerophosphoethanolamine (PEP-), diacylglycerophosphoinositol (PI), diacylglycerophosphoglycerol (PG), sphingomyelin (SM), ceramide (Cer), glucosylceramide (GlcCer), cholesterol derivative (chol der) and cholesterol ester (CE).

**Figure S4.**
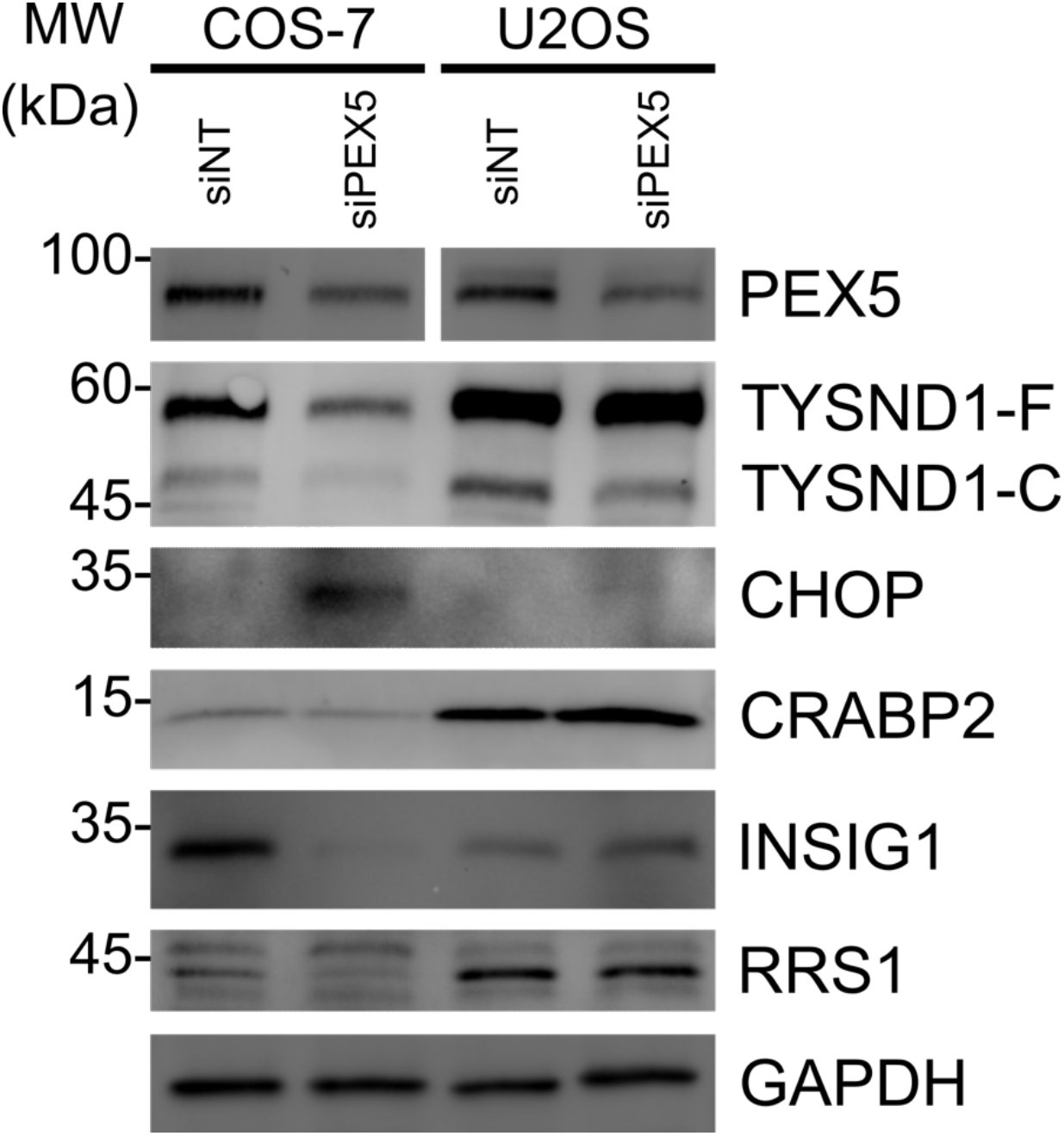
PEX5 silencing reveals common and distinct changes relative to siLONP2. Immunoblotting of non-targeting (*NT*)- and *PEX5*-silenced COS-7 cells and U2OS cells, examining CHOP (ISR), CRABP2 (RA signaling), INSIG1 (cholesterol handling), and RRS1 (ribosomal biogenesis). GAPDH used as a loading control.

### Online supplemental material

Table S1 shows all RNA sequencing data. Table S2 shows all lipidomics data.

## Materials and methods

### Culturing of cell lines

COS-7 (ATCC, CRL-1651) and U2OS (ATCC, HTB-96) cells were cultured in Dulbecco’s modified Eagle medium (DMEM) containing 4.5 gl^-1^ glucose, GLUTA-PLUS and sodium pyruvate (Wisent, 319-027-CL) supplemented with 100 μM non-essential amino acids (Gibco, 11140050) and 10% fetal bovine serum (Wisent, 085-150). All cells were cultured at 37 °C and 5% CO_2_ in a humidified incubator and regularly tested for mycoplasma using the MycoAlert mycoplasma detection kit (Lonza, LT07-418).

### Transfection and RNA interference of cells

For transient transfection, 1 μg plasmid DNA per well was transfected into cells for 16 h at a confluency of 60% in 6-well plates using Lipofectamine 3000 (Invitrogen, L3000001) according to the manufacturer’s recommendations. The following plasmids were used: CFP-SKL (Neuspiel et al., 2008), GFP-LC3 (Kabeya et al., 2000) and pCMV3-hLONP2 (SinoBiological, HG25229-UT). The siRNA-mediated knockdown of proteins was performed for 144 h or 168 h using Lipofectamine RNAiMAX (Invitrogen, 13778150) diluted in Opti-MEM (Invitrogen, 31985-070) and checked by immunoblotting. Briefly, cells with a confluency of 30–40% were transfected with specific siRNAs or a non-targeting control according to the manufacturer’s instructions. After 72 h transfection, the cells were collected, reseeded into 6-well plates, and transfected with the same siRNAs or a non-targeting control for another 72 h or 96 h transfection. The following human siRNAs were used: LONP2 (Dharmacon, ON-TARGETplus, J-005933-06-0005), LONP2 (Dharmacon, Accell, A-005933-14-0005), PEX5 (Dharmacon, ON-TARGETplus, J-015788-07) and non-targeting control (D-001810-10).

### Adenovirus infection

C-terminal YFP-fused PEX3 was subcloned into D2-MCS viral vector (BioVector) under the control of a CMV promoter as described previously (Sugiura et al., 2017). Adenoviral PEX3-YFP was used to infect cells at 100 pfu per cell in the presence of 4 μg/ml polybrene (Sigma, H9268-10G) for 16 hours.

### Immunofluorescence microscopy

Cells were seeded into 24-well plates on glass coverslips and fixed in 5% PFA in PBS, at 37°C for 15 min, then washed three times with PBS, followed by quenching with 50 mM ammonium chloride in PBS. After three washes in PBS, cells were permeabilized in 0.1% Triton X-100 in PBS, followed by three washes in PBS. The cells were blocked with 10% FBS in PBS, followed by incubation in 5% FBS in PBS for 16 h at 4°C with primary antibodies (1:1,000) to the following: LAMP1 (Cell Signaling, 9091), p62 (Cell Signaling, 8025) and PMP70 (Sigma, SAB4200181). After three washes with 5% FBS in PBS, cells were incubated in 5% FBS in PBS for 1 h at RT with appropriate secondary antibodies (1:2,000): cross-adsorbed goat anti–rabbit IgG (H+L) Alexa 594 (Invitrogen, A11012), anti–mouse IgG (H+L) Alexa 647 (Invitrogen, A21235) and anti–rabbit IgG (H+L) Alexa 647 (Invitrogen, A21244). For cholesterol or lipid droplets staining, cells were incubated with 50 μg/ml Filipin (Sigma, SAE0087) for an hour or with 1 μg/ml Nile Red (Sigma, 72485) for 10 minutes, respectively. After five washes in 5% FBS in PBS, coverslips were mounted onto slides using fluorescence mounting medium (Dako). Cells were imaged using a 100× objective NA1.4 on an IX83 inverted microscope (Olympus) with appropriate lasers using a spinning disc system microscope (Yokogawa) coupled to a Neo camera (Andor). For quantification analysis, 30 cells in each condition were randomly chosen and counted based on the definitions on main figures (number and size of peroxisomes: Fig. 1b and 1d; percentage of peroxisomes colocalized with CFP-SKL: Fig. 1f and Fig. S1; number of LC3-p62 puncta on peroxisomes: Fig. 1i; mitochondrial network analysis: Fig. 1j; number of lipid droplets: Fig. 5a; percentage of endolysosomes colocalized with cholesterol: Fig. 5c, d, and f).

### Cell lysates and immunoblotting

To obtain whole cell lysates for western blot analysis, cells were harvested by centrifugation at 15,000 *g* for 10 min and solubilized with HEPES lysis buffer (1% (wt/vol) Triton X-100, 1 mM EDTA, 50 mM HEPES pH 7.4, and 150 mM NaCl). The total protein concentration was determined by using a Bradford assay (Bio-Rad) and the lysates were mixed with sample buffer (10% (vol/vol) glycerol, 2% (wt/vol) SDS, 0.005% (wt/vol) bromophenol blue, 60 mM Tris–HCl pH 6.8 and 10% (vol/vol) β-mercaptoethanol) and heated at 95 °C for 5 min before loading, followed by SDS-PAGE using gradient gels with ranges of 4-16% acrylamide. Proteins were transferred to nitrocellurose membranes and primary antibodies (1:1,000) to the following proteins were used for the immunodetection of proteins using horseradish peroxidase-conjugated anti-mouse (Cytiva, NA931) and anti-rabbit (Cytiva, NA934) secondary antibodies (1:1,000): CHOP (Cell Signaling, 5554), Cleaved Caspase-3 (Cell Signaling, 9661), Cleaved Caspase-7 (Cell Signaling, 9491), CRABP2 (Proteintech, 10225-1-AP), GAPDH (Abcam, AB9485), INSIG1 (Santa Cruz, sc-390504), LC3 (Novus Biological, NB100-2331), LONP1 (Proteintech, 15440-1-AP), NF-κB (Cell Signaling, 6956), p62 (Cell Signaling, 8025), PEX5 (Cell Signaling, 83020), Phospho-eIF2α (Sigma, SAB4504388), Phospho-γH2AX (Abcam, AB2893), PMP70 (Sigma, SAB4200181), Puromycin (Sigma, MABE343), RRS1 (Proteintech, 15329-1-AP), Thiolase (Sigma, HPA007244), TYSND1 (Sigma, HPA030304), and Vinculin (Sigma, V4505).

### RNAseq analysis

Total RNA was extracted from each cell using RNeasy Plus (QIAGEN) after 168 h siRNA transfection, and the quantity and quality was determined using Bioanalyzer (Agilent Technologies). Single read sequencing was performed with NextSeq 500 (Illumina) using 75 cycles High Output kit v2. Sequences were trimmed for sequencing adapters and low quality 3’ bases using Trimmomatic version 0.35 (Bolger et al., 2014) and aligned to the reference *Chlorocebus sabaeus* genome version ChlSab1.1 (gene annotation from Ensembl 104) or *Homo sapiens* genome version GRCh38 (gene annotation from Gencode version 37) using STAR version 2.7.1a (Dobin et al., 2013).

Raw gene expressions counts were obtained directly from STAR, and normalization and differential expression analysis were performed using DESeq2 version 1.30.1 (Love et al., 2014). Metascape software was used for gene ontology analysis (Zhou et al., 2019). We used unadjusted *p*-values for U2OS analysis as false discovery rate calculation has low power in datasets with small effect sizes (i.e. low fraction of changed genes in RNAseq) (Krzywinski and Altman, 2014), and in our dataset Benjamini-Hochberg procedure resulted in over-represented *p*.adjusted values (Table S1). For comparison analyses between COS-7 and U2OS, gene identifiers were converted from *C. sabaeus* to *H. sapiens* using the R/Bioconductor package biomaRt (version 2.48.3). The data are available in NCBI GEO repository (accession no. GSE224992).

### RT-qPCR

Total RNA was isolated from each cell using RNeasy kit (QIAGEN) after 168 h siRNA transfection. RT–qPCR and data analysis were performed at IRIC Genomics Platform (University of Montreal). All reactions were run in triplicate and the average values of Ct were used for quantification. The relative quantification of target genes was determined using the ΔΔCT method. Primers for target genes are as follows: LONP2 (COS-7) (Fw: gtgggaagatcagtggccaa, Rv: gcctgtgtcctcgaatgtca), LONP2 (U2OS) (Fw: ggggacgtgatgaaggagtc, Rv: agcattggtcagctggtact), TGM2 (Fw: cggatgctgtgtacctggac, Rv: cttgatgaacttggccgagc), TBP (endogenous control) (Fw: gaacatcatggatcagaacaaca, Rv: atagggattccgggagtcat) and YWHAZ (endogenous control) (Fw: gcaattactgagagacaacttgaca, Rv: tggaaggccggttaatttt)

### Puromycylation assay

Translation intensity was monitored by the incorporation of puromycin as previously described (Schmidt et al., 2009). After 7 days siRNA transfection, cells were pulsed with 10 μg/ml puromycin (Sigma, P8833) at 37°C for 10 minutes. Cells were then harvested as described above and blots probed with anti-puromycin antibodies, as described above.

### Retinoic acid stimulation

COS-7 cells were incubated in presence or absence of 5 μM retinoic acid (Sigma, R2625) at 37°C for 24 h before isolating total RNA for analysis by qRT-PCR.

### Untargeted Lipidomics

Lipid extraction, sample and data analysis were performed using a previously validated semi-quantitative untargeted lipidomic workflow (Forest et al., 2018). In brief, lipids were extracted from each cell after 168 h siRNA transfection and spiked with six internal standards: LPC 13:0, PC19:0/19:0, PC14:0/14:0, PS12:0/12:0, PG15:0/15:0, and PE17:0/17:0 (Avanti Polar Lipids). Protein concentration was determined using a colorimetric-based assay based on the Bradford dye-binding method and samples were injected into a 1290 Infinity high resolution HPLC coupled with a 6530 Accurate Mass quadrupole time-of-flight (LC-QTOF) (Agilent Technologies) via a dual electrospray ionization (ESI) source. Elution of lipids was assessed on a Zorbax Eclipse plus column (Agilent Technologies) maintained at 40 °C using an 83 min chromatographic gradient of solvent A (0.2% formic acid and 10 mM ammonium formate in water) and B (0.2% formic acid and 5 mM ammonium formate in methanol/acetonitrile/methyl tert-butyl ether [MTBE], 55:35:10 [v/v/v]). Data acquisition was performed in positive ionisation mode. All samples were processed and analyzed as a single batch. MS quality controls (QCs) were performed by (i) injecting 3 “in-house” QC samples and blanks at the beginning, middle and end of the run and (ii) monitoring six internal standards spiked in samples for signal intensity, mass mass-to-charge ratios (m/z) and retention time (RT) accuracies. Mass spectrometry (MS) raw data processing was achieved as previously described using Mass Hunter B.06.00 (Agilent Technologies) for peak picking and an in-house bioinformatic script [2018, Forest et al] in both Perl and R languages that we developed for: (i) MS feature peak alignment and retention time (RT) correction; (ii) filter of presence: features retained must be present in 80% of samples from at least one group, thereby setting a maximum of 20 missing values within a group; (iii) normalization of signal intensities using cyclic loess algorithm; (iv) imputation of missing values to 90% of lower values. The resulting final dataset included 2,094 high-quality MS signals, thereafter referred to as features, defined by their m/z, RT and signal intensity (listed in Table S2). Statistical analyses are performed using the final dataset on log2-transformed signal intensity data. An unsupervised principal component analysis was first applied followed by independent testing of each feature using or 2-tailed unpaired Student’s *t* test with Benjamini-Hochberg correction for the indicated group comparison. Given the untargeted nature of our lipidomic analysis, only lipid features of interest have been annotated to unique lipids, a step that is crucial given that about 50% of all MS features are duplicate ions of the same lipid. For this study, lipid annotation was focused on the features that passed the selected threshold of significance, namely *P*_*corr*_-value (*P* value after Benjamini-Hochberg multiple testing correction) of < 0.05 and an absolute fold change (FC) |(FC)|> 1.5 (corresponding to |log2 (FC)|> 0.58) for the comparisons siLONP2 vs. siNT in COS-7 and U2OS. Lipid annotation was achieved as previously described in details (Forest et al.) by (i) MS/MS analysis for the 100 most significant lipid features in each comparison and (ii) through data alignment with our in-house database, which contains >500 unique lipids with previously determined MS/MS spectra for the remaining significant features.

### Cholesterol flux analysis

Cholesterol was delivered to cells by a cholesterol-cyclodextrin complex (Sigma, C4951) that contained approximately 40 mg of cholesterol per gram of material. All treatment concentrations were calculated based on cholesterol weight. After 168 h siRNA transfection, cells were incubated with 200 μM cholesterol in culture medium at 37°C for an hour, washed and incubated in culture medium for an hour, followed by fixation and immunofluorescence microscopy.

### Transmission electron microscopy

Electron microscopy (EM) was performed as previously described (Neuspiel et al., 2008). After 168 h siRNA transfection, cells were fixed with 5% PFA and 1.6% glutaraldehyde (GA) in 0.1 M sodium cacodylate buffer (pH 7.4) for 10 min at RT, then further fixed at 4 °C in the same buffer overnight. After washing with 0.1 M cacodylate buffer, cells were fixed with 1% osmium tetroxide for 60 min at 4 °C, washed with water, stained with saturated aqueous uranyl acetate for 45 min at RT, and then gradually dehydrated with a series of increasing concentrations of ethanol (70–100%). After dehydrating with 100% acetone, cells were gradually embedded in Spurr’s resin, and polymerized for 48 h at 60 °C. Samples were sectioned to a 100-nm thickness and sections were mounted on 200-mesh copper grids, and sections were imaged at 120 kV using a FEI Tecnai 12 TEM outfitted with an AMT XR80C CCD Camera System, housed in the Facility for Electron Microscopy Research (FEMR) at McGill University.

### Statistics and reproducibility

Statistical significance was tested by the unpaired two-tailed Student’s *t* test using the GraphPad Prism software (version 8.4.3). The values of all individual cells and the mean of each experiment are shown for the quantification analysis. Errors bars displayed on graphs represent the means ± SD of at least three independent biological replicates. Graphs were generated using Prism or R.

